# Timing the regional spread of PRRSV-2 variants across the United States

**DOI:** 10.64898/2026.03.12.711334

**Authors:** Joao P. Herrera da Silva, Igor Paploski, Mariana Kikuti, Nakarin Pamornchainavakul, Cesar A. Corzo, Kimberly VanderWaal

## Abstract

Porcine Reproductive and Respiratory Syndrome Virus 2 (PRRSV-2) represents a major threat to the global swine industry. The epidemiological dynamics of PRRSV-2 are characterized by the recurrent annual emergence of dozens of variants. Long-distance spread of PRRSV-2 is largely driven by animal shipments. Spatiotemporal dynamics of PRRSV-2 in the USA have been explored; however, how fast variants spread to new regions after their emergence remains unclear, and this information could improve preparedness. To address this, we analyzed 14,835 sequences, retrieved from the Morrison Swine Health Monitoring Project (MSHMP), representing 156 variants sampled from 2015 to 2024, covering the five major swine-producing regions in the USA: the Upper Midwest (UM), Lower Midwest (LM), Atlantic Seaboard (AS), Northeast (NE), and Great Plains (GP). Time to spread was assessed using the time-to-dispersal event analysis and waiting time analyses. Genetic diversity was measured using Hill numbers. The UM had the highest variant richness (n=123), followed by the LM (n=47), AS (n=35), NE (n=45), and GP (n=38). Of the 62 variants that initially emerged in the UM, 17 later spread to other regions. The UM also received the highest number of variant introductions (n=24), followed by LM (n=14), NE (n=14), AS (n=4), and GP (n=7), highlighting regional differences in connectivity and risk. Our results suggest faster dispersal corridors among interior regions (e.g., GP to UM and LM to UM, ∼1.2–2.0 years) and slower for coast to interior pathways (AS to interior, ∼2–3 years). These findings may help anticipate the risk of PRRSV-2 variant introduction and provide more accurate dispersal time estimates, which are useful for improving epidemiological models and disease preparedness.

## Introduction

It is estimated that around 50% of the diseases that have emerged during the 21st century, affecting plants, animals, and humans, are caused by viral agents [1–4]. Disease emergence can be defined as the first report of a new disease or a genetically distinct strain of a known pathogen, an increase in incidence, a spillover to a new host, or the spread into geographical regions where it had not been previously reported [2]. Numerous factors are associated with disease emergence, including high mutation rates, recombination events, genetic bottlenecks, selective pressures, land-use changes, and global connectivity [1, 5–15]. The rapid diversification of some viral species generates heterogeneous populations composed of multiple genetic variants that coexist in the environment [16–19]. This diversity can confer robustness to the population, understood as the ability to preserve functionality in the face of genetic or environmental perturbations, thereby increasing the resilience of viral population [20–22]. In practice, this complex dynamic poses challenges to the durability of control strategies, requiring constant updates of vaccines and resistance genes, which are occasionally overcome by highly diversifiable pathogens [23–26]. Among viruses that threaten livestock health, Porcine Reproductive and Respiratory Syndrome Virus type 2 (PRRSV-2) stands out as a prominent example. PRRSV-2 exhibits rapid genetic diversification, and one of the main factors contributing to strain emergence in new areas has been the movement of animals [27–30].

*Betaarterivirus americense* (Porcine Respiratory and Reproductive Syndrome Virus-2/PRRSV-2) is an enveloped, positive-sense, single-stranded RNA virus that belongs to the *Arteriviridae* family and order *Nidovirales* [31]. PRRSV-2 exhibits rapid evolutionary dynamics, driven by high mutation rates and frequent recombination events [27–29]. Based on the phylogenetic relationships of the ORF5 gene, PRRSV-2 is hierarchically classified according to genetic variants, which are nested within phylogenetic sub-lineages and lineages [18, 32–34]. Typically, viruses belonging to the same PRRSV-2 genetic variant share an average genetic distance of approximately 2.5% and are ∼5% different from the most closely related variant [18]. PRRSV-2 populations in USA display complex dynamics marked by the recurrent emergence of new genetic variants. It is estimated that over the last decade, an average of 18 variants has emerged annually in the USA, with at least 48 co-circulating in the country each year [18, 33].

Clinical signs of PRRSV infection include reproductive failure in sows, which can manifest as abortion, stillborn, mummification, respiratory issues such as pneumonia, and also reduced weight gain [35, 36]. Since its emergence almost four decades ago, PRRSV-2 has posed a significant threat to the swine industry in the world [37–40]. Estimates suggest PRRSV-2 causes over $1.2 billion in losses annually in the USA alone [41]. PRRSV-2 is also the most sequenced non-human virus worldwide, with sequences primarily generated by veterinarians and producers through routine diagnostic testing [18].

PRRSV-2 can be transmitted both horizontally and vertically. Vertical transmission can occur through semen or trans-uterine transmission [42–45]. Horizontal transmission occurs through either direct contact between animals or via fomites, as well as through airborne spread [30, 46–48]. Dispersal can occur over short distances via airborne spread or accidentally through management practices [49]. In the United States, the swine industry is predominantly organized as a multi-site production system, which results in intense movement of animals both within and across states. Approximately 71% of pigs are finished in locations different from where they were farrowed [55]. While much of this movement occurs within states, a substantial fraction involves interstate transportation [55], and animal shipments have been identified as a factor strongly associated with the long-distance spread of PRRSV-2, particularly through the movement of pigs from nurseries to finishing sites [34, 50–54].

Phylogeographic analyses integrating sequence data from North America have revealed the main geographic pathways of PRRSV-2 dissemination [56, 57], with asymmetric patterns of dispersal among regions identified. The U.S. Upper Midwest has been consistently indicated as one of the primary sources of PRRSV-2 spread to other regions, particularly to the Northeast and the Lower Midwest, while also plays a role as a recipient region [56, 57]. In contrast, the Atlantic Seaboard has been identified as playing a source role, contributing to the dissemination of PRRSV-2 mainly toward the Upper Midwest and the Northeast [56, 57]. However, these phylogeographic models were performed at the sub-lineage level, focusing on the diverse and prevalent Lineage 1 [33, 56], which has pairwise genetic distances ranging from 10.2% to 14.2% [32]. This broad diversity limits our ability to resolve recent variant emergence and spread events, given that spatiotemporal dynamics of sub-lineages occur over the course of decades. In addition, sub-lineages are usually too large and too diverse to capture the dynamics of specific outbreak events, which are usually attributed to a single variant [58].

Thus, although regional transmission pathways of PRRSV-2 have already been characterized at the sub-lineage level, open questions remain regarding how rapidly dispersal occurs following the emergence of new genetic variants. However, the recently adopted variant-based classification framework provides higher resolution for tracking the emergence of new variants and enabling the reconstruction of epidemiological links at finer spatial and temporal scales [18]. The epidemiological dynamics of PRRSV-2 variants are marked by frequent emergence of new variants and displacement of others. Here, we use PRRSV-2 genetic variants to quantify the diversity of the co-circulating variants at the regional level, identify regions that frequently exchange variants, and estimate time-lags between their emergence in a given region and their subsequent dissemination to other regions.

This information has the potential to add an additional layer to our understanding of the real risks associated with PRRSV-2 variant emergence, dispersal, and expansion into new regions by placing these processes on a temporal scale. This framework may enable intensified surveillance in highly connected areas that function as rapid dissemination corridors and support the implementation of risk-based alert systems when highly pathogenic or highly transmissible variants emerge. These alert systems may also support preparedness for potential new introductions by prompting intensified diagnostic testing and providing a unique opportunity for coordinated information sharing among production systems, helping to contain the spread of variants of concern.

## Methods

### Data

A total of 35,167 ORF5 sequences were obtained from Morison Swine Health Monitoring Project (MSHMP), which includes sequences generated from routine diagnostics from breeding and growing pig sites of its participants, representing more than 60% of U.S. breeding population [59]. Sequences collected before 2015 were removed from the data set to avoid misclassification, since variant classification was established using only sequences from 2015 onwards [18, 60]. In total 27,388 sequences collected after 2015 were classified into variant using *PRRSLoom*-Variants web tool (update September 2025) [18]. In the variant classification system, variants are defined based on phylogenetic relationships and genetic distance; sequences belonging to the same variant have an average genetic distance of approximately 2.5% [18]. Overall, our dataset comprised an average of 145.8 sequences per variant, with a median of 24.5. Since we aimed to determine the time of variant spread, sequences assigned to an undetermined sub-lineage or unclassified variant were removed; such sequences could not be matched to a well-resolved phylogenetic clade. Sequence belonging to vaccine-like variants (variant 5A.1: Ingelvac PRRSV MLV–GenBank ID AF066183.4; variant 7.1: Prime Pac PRRSV RR–GenBank ID DQ779791.1; variant 8A.1: Ingelvac PRRSV ATP–GenBank ID DQ988080.1; variant 8C.1: Fostera PRRSV–GenBank ID AF494042), were removed. After applying all filters, 14,835 sequences were retained and has used for the following analyses.

### PRRSV-2 diversity in different regions

The United States can be divided into six swine-producing regions according to the Swine Health Information Center’s (SHIC) Rapid Response Program [61]. For the purposes of this study, we adopted the same regional subdivision used by Makau *et al.* [62], which is based on the SHIC classification but with a slight modification: the Midwest is further divided into two subregions: Upper Midwest and Lower Midwest. In this analysis, we focused on the five regions with the highest densities of swine production in the United States: Upper Midwest, Lower Midwest, Atlantic Seaboard, Great Plains, and Northeast (Supplementary Figure S1), as these were also the regions for which we had access to the largest number of sequences. To assess diversity of PRRSV-2 variants across different regions, we estimated the Hill numbers (the effective number of variants) with scaling parameter *q* ranging from 0 to 2 [63] using the iNEXT package implemented in R [64]. In summary *q* = 0 represents the variant richness (i.e., absolute number of distinct variants detected), counting all variants equally without weighting by relative abundance, and thus emphasizing rare variants equally to prevalent ones; *q* =1 accounts for evenness, by weighting variants according to their relative abundance in the region, corresponding to the exponential of Shannon entropy; and q = 2 gives more weight to dominant variant, which corresponds to the inverse Simpson Index [65, 66]. Due the imbalance of the number of sequences in different regions, we evaluated whether the sampling effort (i.e., number of sequences) was sufficient to capture the full diversity in their respective regions through the construction of variant accumulative curves for each region using iNEXT, applying both rarefaction and extrapolation approaches [64].

### Estimating time-lags between the first emergence of a variant in their region of origin to dissemination to other regions using time-to-dispersal event analyses

To better understand the relative contribution of local emergence and importation to the composition of PRRSV-2 variant diversity in each region, and to further assess the time interval between the local emergence of variants in one region and their subsequent dissemination to others, we estimated for each region: (i) the number of variants that emerged locally; (ii) the number of variants imported from other areas into the region; (iii) the number of variants exported from the original region to recipient regions; and (iv) the number of variants that apparently remained restricted to the region of origin. Based on uncensored observations, we also estimated the interval (in years) between emergence in the region of origin and detection in a recipient region. Emergence events were defined in this study as the detection of at least five sequences of a given variant within a specific region. The date on which a variant reached this threshold was considered its emergence date. For the purpose of this analysis, variants that did not reach a minimum of five sequences within their respective regions, hereafter referred to as underrepresented variants, were excluded due to their limited epidemiological information. The threshold of five sequences was chosen to minimize the influence of isolated detections and variants showing minimal evidence of a sustained circulation were considered.

For the time-to-dispersal event analyses, we applied an approach analogous to the *Mainland model*, assuming that a single region serves as the source for pathogen dispersal to other regions. For example, the time to dispersal between Region A → B and A → C were independently assessed; the potential for variant movement through intermediate regions was not accounted for, e.g. A→B→C. Thus, this approach captures time-lags between emergence in one region and subsequent detection in each other region, but does not directly assess the route of spread. Only variants that reached at least five sequences in a given region were considered.

To account for whether emerging variants detected in the most recent years of surveillance had sufficient time to disperse to other regions (i.e., to define right censoring), we introduced a hypothetical extinction time, not intended to represent the true biological disappearance of a variant, but based on the average interval between the first and last detection of each variant, which was approximately three years. Accordingly, variants that emerged within the final three years of sampling and were not detected outside their region of origin were treated as right-censored due to insufficient follow-up time for dispersal. Conversely, variants that emerged earlier than this three-year window and likewise did not disperse beyond their region of origin were treated as left-censored at the time of their last detection, reflecting sufficient opportunity for dispersal without evidence of spread. In summary, variants emerging within the last three years without evidence of dispersal were right-censored, whereas earlier non-dispersing variants were left-censored. Dispersal time was defined as the interval between the fifth detection of a variant in the region of origin and the fifth detection in each recipient region. The time to dispersal event was estimated in R version 3.6.0+ [67] using the *survival* package [68]. The Kaplan–Meier estimator was implemented using the *survfit2* function. The time-to-dispersal event curves were generated for each recipient region, with time defined as the interval (in years) between the emergence and dispersal of each variant, and censoring status encoded accordingly. Median time-to-dispersal event and 95% confidence intervals were extracted from the fitted survival objects using the *surv_median* function. Differences in the time-to-dispersal event distributions among recipient regions were evaluated with the log-rank test, as implemented in the *survdiff* function from the survival package.

### Estimating the time-lag between variant emergence/introduction and subsequent dispersal to another region using phylogeographic analysis

We also estimated dispersal times using a phylogeographic approach, implemented with waiting time analyses. Waiting time is defined as the time lag between variant emergence/introduction and subsequent dispersal to another region. The analysis was performed separately for each variant. To minimize uncertainty and stochastic effects in these analyses, we restricted the dataset to variants represented by more than 100 sequences in total and detected in at least two regions, thereby ensuring sufficient temporal depth and geographic representativeness. In total, fifteen variants were included.

Multiple sequence alignments of the ORF5 gene were performed separately for each variant using MAFFT v.7 algorithm [69] with default parameters. To assess the presence of a temporal signal, a root-to-tip regression analysis between genetic distance and sampling date in TempEst [70] was performed. The best nucleotide model was determined using the ModelTest-NG [71] based on Bayesian Information Criterion (BIC); the GTR+I+G4 was selected as the best-fit model for all datasets. The phylogenetic analyses were performed in RAxML-NG using 1,000 bootstrap replicates [72]. Time-scaled trees were inferred using BEAST 1.10.4 [73], applying the General Time Reversible (GTR) substitution model with empirical base frequencies and a Gamma distribution plus invariant sites (GTR+I+G). An uncorrelated log-normal relaxed molecular clock was used, along with a Skygrid coalescent model with 50 transition-points for the change in population size [74]. The spatial-temporal histories for each variant were reconstructed using a discrete-state continuous-time approach with the Bayesian Stochastic Search Variable Selection approach, with swine-producing regions (Supplementary Figure S1) used as the discrete-trait variable [75]. Posterior probability distributions were estimated by running a Markov chain Monte Carlo (MCMC) for 300 million iterations, with samples collected every 30,000 states. The convergence of the MCMC chains was determined based on effective sample size of at least 200, and by the degree of interdependence of the samples assessed based on the degree of mixing of the parameters, assessed using Tracer v.1.7.2 [76]. 9,000 posterior trees were kept for the following analysis discarding the initial 10% of the chains as burn-in. The waiting times of each variant in their respective geographic regions were estimated as the average time required for a given variant sampled in a particular region to coalesce in the previous generation in a different region. Waiting times were determined using Posterior Analysis of Coalescent Trees (PACT) v0.0.5 [77].

## Results

### PRRSV Diversity and Variability in different USA regions

We assessed the diversity of PRRSV-2 variants across the five major swine-producing regions in the United States: Upper Midwest, Lower Midwest, Atlantic Seaboard, Great Plains, and Northeast. The South and Western regions were excluded due to the limited number of available sequences and their relatively minor contribution to national swine production.

Variant diversity was quantified using Hill numbers at three orders (q = 0, 1, and 2), representing variant richness, abundance of common variants, and dominance of variants, respectively. The Upper Midwest region exhibited the highest variant richness (q = 0), with a total of 123 distinct variants, followed by the Lower Midwest (47 variants), Northeast (45 variants), Great Plains (38 variants), and Atlantic Seaboard (35 variants) (Figure 1, Supplementary Figure 2 and Supplementary Table 2). In comparison with the other regions, the Lower Midwest region exhibited the highest evenness according to the Hill number of order q=1, followed by the Northeast and Upper Midwest, which showed similar values, then the Atlantic Seaboard, and finally the Great Plains. These results indicate that the Lower Midwest is composed of a relatively large number of variants with more balanced relative abundances, when compared to the other regions (Figure 3). When examining Hill numbers of order q=2, which emphasize dominant variants, the Lower Midwest again showed the highest diversity, suggesting the presence of several highly prevalent variants. This was followed by the Atlantic Seaboard and Northeast, then the Great Plains, and lastly the Upper Midwest, which showed the lowest diversity under this metric (Figure 1).

**Figure 1.**
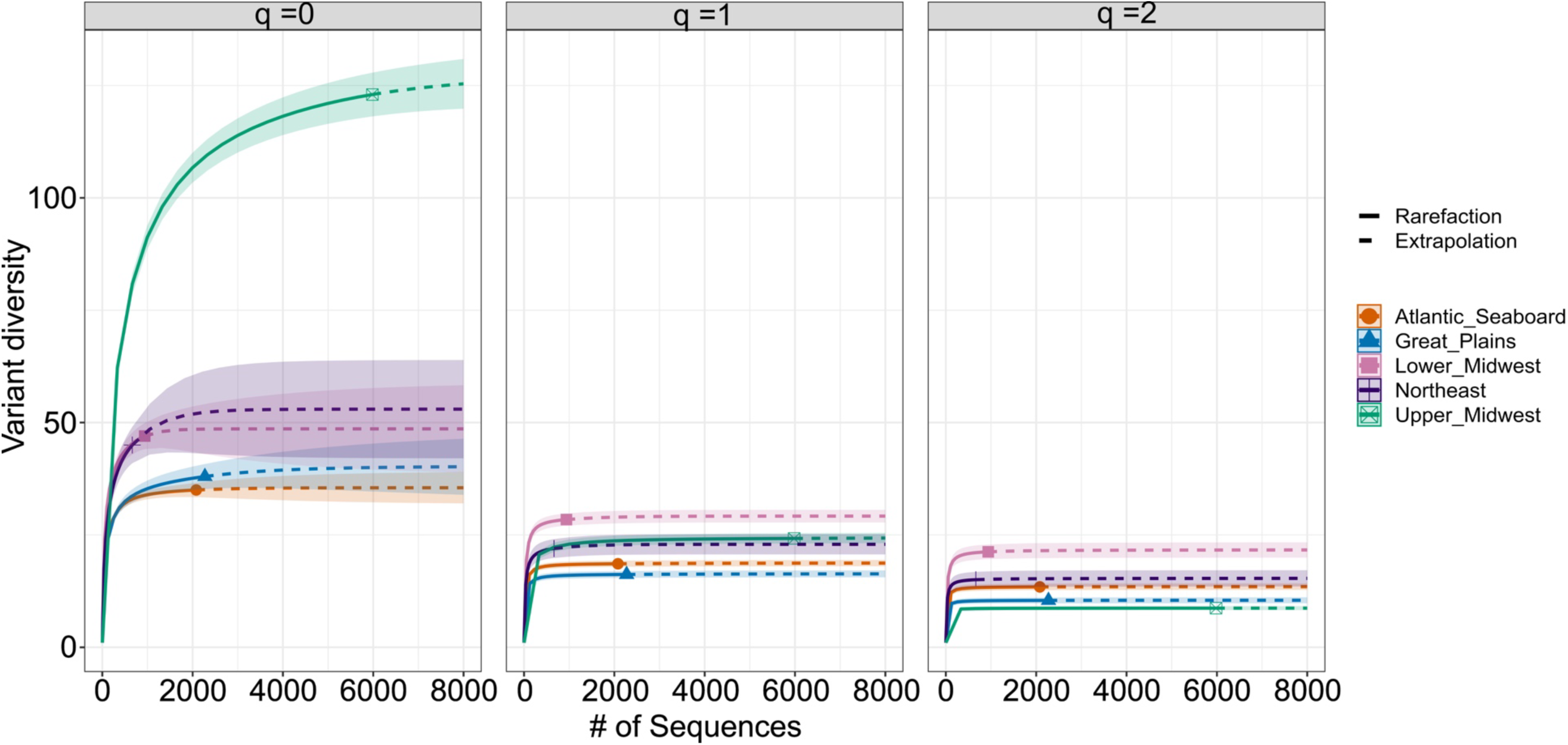
Variant diversity profiles of PRRSV-2 across USA swine-producing regions. (A) Number of variants detected per region. (B) Number of sequences obtained per region. (C) Rarefaction and extrapolation curves of Hill numbers for orders q = 0 (richness), q = 1 (Shannon diversity), and q = 2 (Simpson diversity). Solid lines indicate rarefaction and dashed lines extrapolation, with shaded areas representing 95% confidence intervals. Colors correspond to swine-producing regions: Atlantic Seaboard (orange), Great Plains (blue), Lower Midwest (purple), Northeast (pink), and Upper Midwest (green).

**Figure 2.**
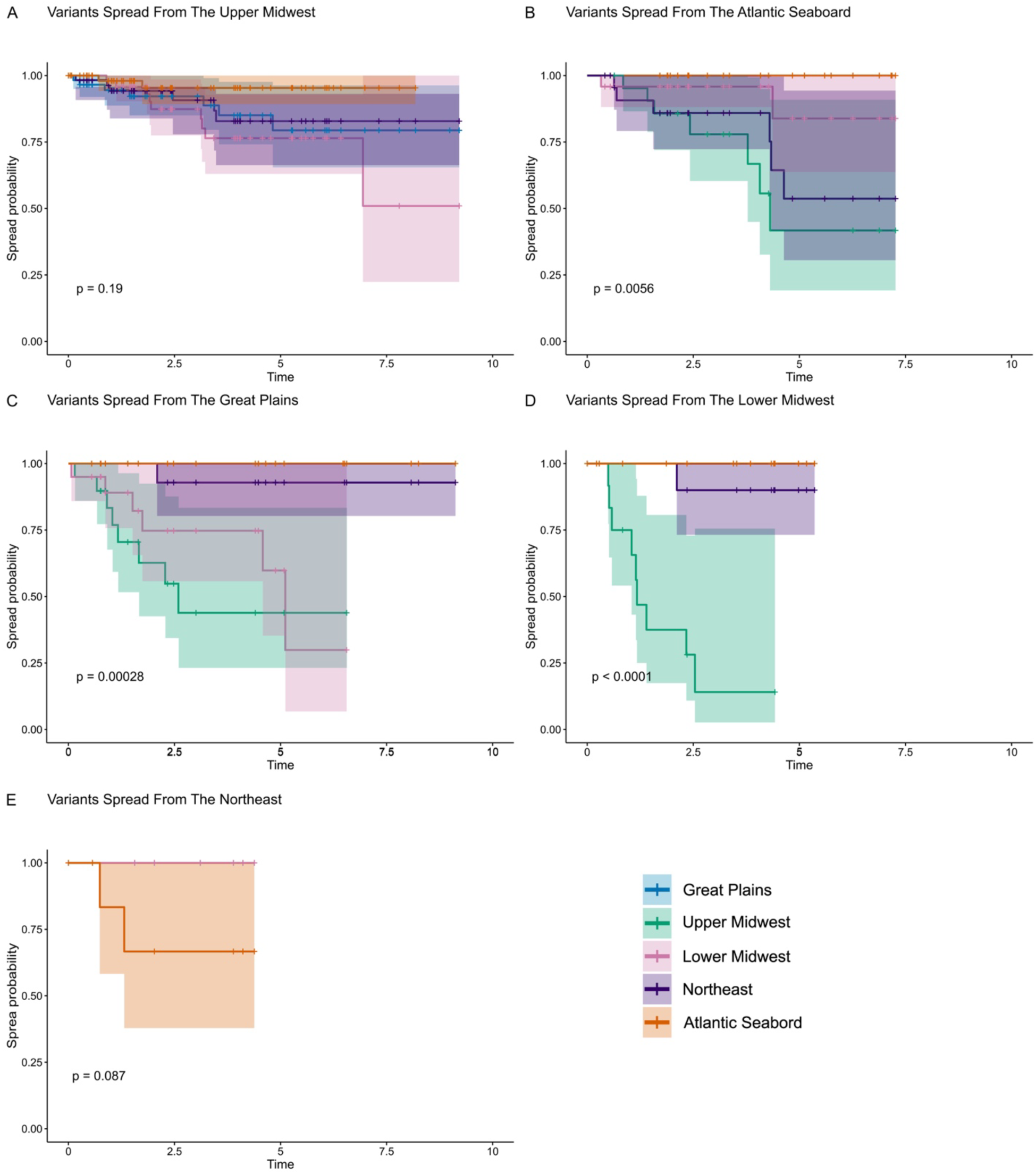
Kaplan–Meier the time to dispersal event curves showing the time between local emergence of PRRSV-2 variants in the original region and their first detection in recipient regions. Panels A–E correspond to variants emerging in each of the five major swine-producing regions: Upper Midwest (A), Atlantic Seaboard (B), Great Plains (C), Lower Midwest (D), and Northeast (E). The y-axis represents the probability that a variant has not yet been detected outside its region of origin at a given time (x-axis, in years). Colored lines represent different recipient regions, and shaded areas correspond to 95% confidence intervals. The log-rank p-values are shown for comparisons across recipient regions. Note that the time to dispersal event probabilities decrease as variants are detected in new regions over time. Censored observations are indicated by “+” symbols along the time to dispersal event curves.

**Figure 3.**
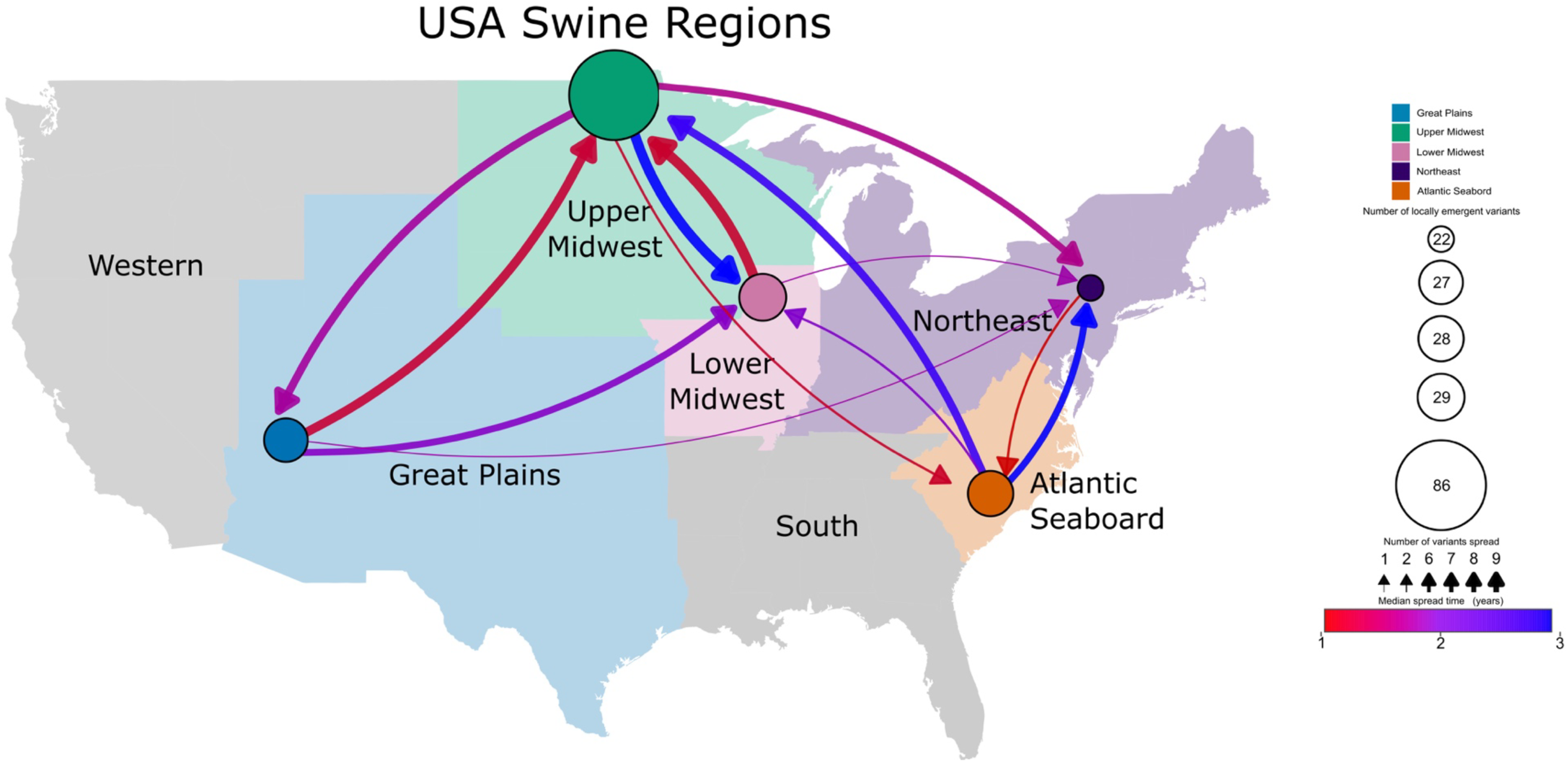
Network of PRRSV-2 variant dispersal among USA regions. Each node represents a geographic region, and its size is proportional to the number of locally emergent variants detected within that region. Arrows depict directional dispersal events of variants between regions; arrow width indicates the number of variants spread, and arrow color corresponds to the median spread time (in years), as shown in the color bar. Warm colors represent faster and cool colors indicate slower spread dynamics. Regional colors correspond to the categories shown in the legend.

In the Upper Midwest, variant 1C.5 was predominant, accounting for nearly 30% of all sequences analyzed from this region across 2015-2025 (Supplementary Figure 2 and Supplementary Table 2). Among the 123 variants identified in the Upper Midwest, four collectively represented more than 50% of the sequences (Supplementary Figure 2 and Supplementary Table 2). In contrast, 47 variants were detected in the Lower Midwest; the predominant variant was 1C.2, which accounted for 11% of all sequences from this region. Of the 47 variants detected there, eight together comprised approximately 50% of all sequences (Supplementary Figure 2 and Supplementary Table 2). In the Northeast, the predominant variant was 1A.28, representing 14% of the sequences. Out of 45 variants identified in this region, seven accounted collectively for about 50% of the sequences (Supplementary Figure 2 and Supplementary Table 2). In the Great Plains, variant 1E.1 was the most prevalent, accounting for 21% of regional sequences. Among the 38 detected variants, five comprise 50% of all sampled sequences, with most belonging to sub-lineage 1H (Supplementary Figure 2 and Supplementary Table 2). Finally, in the Atlantic Seaboard, the most frequently detected variant was 1C.1, which represented 16% of all sequences though it has not been detected since 2019. Sub-lineage 1A appears most frequently in this region, although the majority of these sequences are assigned as 1A-unclassified variants. In this region, 35 variants were reported, of which seven collectively accounted for 50% of the sequences analyzed (Supplementary Figure 2 and Supplementary Table 2). Together, these findings suggest that although the Upper Midwest harbors the greatest number of PRRSV-2 variants (as shown by q=0), most of these variants occur at low frequencies, with only a few being highly abundant. In contrast, the Lower Midwest is characterized by a more even distribution of variants.

Although the number of sequences varied across regions, we constructed rarefaction curves for the three Hill numbers to evaluate whether sampling intensity was sufficient to capture regional diversity. None of the rarefaction curves reached a saturation plateau for the richness estimates (q=0). However, the rarefaction curves reached a plateau for abundance (q=1) and dominance (q=2, Figure 1). These results suggests that additional rare variants could still be detected with increased sampling effort, but the overall abundance and dominance likely was captured by our sampling effort.

### Regional emergence, dissemination, and temporal lags reveal connectivity patterns between swine regions in USA

Regions that functioned as centers of emergence, sources, or recipients of PRRSV-2 variants were characterized based on patterns of local emergence, importation, exportation, and estimated interregional dispersal times. The time interval between the fifth detection in the region of origin and the fifth detection in the recipient region was used to estimate dispersal times.

After removing underrepresented variants, defined here as those that did not reach a minimum level of sustained circulation (at least 5 sequences detected), we observed clear regional variation in the number of variants that emerged locally, remained geographically restricted, or disseminated to new regions, as well as in the level of variant importation and time elapsed between emergence and inter-regional dispersal (Tables 1–3; Figures 2–3). Together, these patterns reveal marked asymmetries in connectivity within the PRRSV-2 variant dispersal network.

**Table 1:** Summary of the total number of variants detected in each region, the number of variants that emerged locally, the number of variants that remained restricted to their region of origin, the number of locally emerged variants that were also detected in other regions, and the number of variants introduced from other regions. \**Underrepresented variants* are variants that did not reach a minimum of five sequences within their respective regions hereafter referred.

| Original region | Total variants | After excluding *underrepresented variants* * |  |  |  |  |
| --- | --- | --- | --- | --- | --- | --- |
|  |  | Total variants | Locally emerged | Regionally restricted | Spread into other regions | Introduced from other regions |
| Atlantic Seaboard | 35 | 28 | 24 | 15 | 9 | 4 |
| Great Plains | 38 | 27 | 20 | 12 | 8 | 7 |
| Lower Midwest | 47 | 29 | 15 | 6 | 9 | 14 |
| Northeast | 45 | 22 | 8 | 6 | 2 | 14 |
| Upper Midwest | 123 | 86 | 62 | 45 | 17 | 24 |

**Table 2:** Number of variants that spread from original regions to their respective recipient regions. Note that the values presented should not be summed across regions, as they are not mutually exclusive: a single variant may be counted in multiple recipient regions if it was exported from its region of origin to more than one destination.

| Number of variants that spread from original regions to each recipient regions |  |  |  |  |  |
| --- | --- | --- | --- | --- | --- |
| Original region | Recipient region |  |  |  |  |
|  | Atlantic Seaboard | Great Plains | Lower Midwest | Northeast | Upper Midwest |
| Atlantic Seaboard | - | 0 | 2 | 6 | 7 |
| Great Plains | 0 | - | 6 | 1 | 8 |
| Lower Midwest | 0 | 0 | - | 1 | 9 |
| Northeast | 2 | 0 | 0 | - | 0 |
| Upper Midwest | 2 | 7 | 9 | 6 | - |

**Table 3.** Average time (in years) between the local emergence of PRRSV-2 variants in the original region and their emergence in each recipient region. n represents the number of variants spread events detected between the source and recipient regions. The Table also reports the average and median time intervals for inter-regional dissemination, along with the corresponding standard deviation. “NA” indicates that the standard deviation could not be calculated due to a single observation.

| Time to Dispersal (in years) |  |  |  |  |  |
| --- | --- | --- | --- | --- | --- |
| Origin Region | Recipient Region | N (number variants) | Average | Median | Standard Deviation |
| Great Plains | Upper Midwest | 8 | 1.31 | 1.10 | 0.82 |
| Great Plains | Lower Midwest | 6 | 2.31 | 1.63 | 2.05 |
| Great Plains | Northeast | 1 | 2.09 | 2.09 | NA |
| Great Plains | Overall | 8 | 1.76 | 1.51 | 1.45 |
| Upper Midwest | Great Plains | 7 | 2.03 | 1.44 | 1.82 |
| Upper Midwest | Lower Midwest | 9 | 2.69 | 1.95 | 1.81 |
| Upper Midwest | Northeast | 6 | 1.90 | 1.72 | 1.41 |
| Upper Midwest | Atlantic Seaboard | 2 | 1.22 | 1.22 | 0.73 |
| Upper Midwest | Overall | 17 | 2.18 | 1.88 | 1.63 |
| Lower Midwest | Upper Midwest | 9 | 1.25 | 1.15 | 0.75 |
| Lower Midwest | Northeast | 1 | 2.11 | 2.11 | NA |
| Lower Midwest | Overall | 9 | 1.33 | 1.16 | 0.76 |
| Northeast | Atlantic Seaboard | 2 | 1.03 | 1.03 | 0.41 |
| Northeast | Overall | 2 | 1.03 | 1.03 | 0.41 |
| Atlantic Seaboard | Upper Midwest | 7 | 2.63 | 2.42 | 1.42 |
| Atlantic Seaboard | Lower Midwest | 2 | 2.35 | 2.35 | 2.86 |
| Atlantic Seaboard | Northeast | 6 | 2.69 | 2.93 | 1.93 |
| Atlantic Seaboard | Overall | 9 | 2.61 | 2.41 | 1.67 |
| Total | Overall | 45 | 2.02 | 1.53 | 1.36 |

The Upper Midwest stands out as the region exhibiting the highest number of locally emerging variants (Table 1; Figure 2A). In addition, it can be characterized as a central hub in the PRRSV-2 dissemination network, functioning both as a major source of spread and as a primary recipient of variants. The Upper Midwest was the only region that exported variants to all other regions, with the Lower Midwest serving as the principal destination (Table 2; Figure 2A and Figure 3). From a recipient perspective, the Upper Midwest received variants from most other regions, with the exception of the Northeast. Despite its high connectivity, the majority of variants that emerged in the Upper Midwest remained geographically restricted to the region (Table 1). Regarding spread dynamics, the Upper Midwest occupied an intermediate position in terms of spread time, with an overall mean time to dispersal of approximately 2.2 years (Table 3). Variants emerging in this region spread more rapidly to the Atlantic Seaboard and Northeast (1.2 – 1.9 years, on average), and more slowly to the Great Plains and Lower Midwest (2.0 – 2.7 years, Table 3; Figure 2A and Figure 3).

The Great Plains exhibited a relatively balanced pattern of variant exchange, receiving and exporting a similar number of variants (Table 1). More than half of 20 locally emerging variants remained geographically restricted (Table 1). The Great Plains disseminated variants to most regions, with the exception of the Atlantic Seaboard; however, all detected variant introductions into the region originated from the Upper Midwest (Table 2). The Upper and Lower Midwest were the primary destinations of variants emerging in the Great Plains. From a time-to-dispersal perspective, the Great Plains occupied an intermediate position, with variants emerging in this region taking on average approximately 1.8 years to be detected in other regions (Table 3; Figures 2C and Figures 3). Notably, dissemination from the Great Plains occurred more rapidly toward the Upper Midwest (1.3 years, on average), consistent with strong connectivity between interior regions (Table 3; Figures 2C and Figures 3).

The number of locally emerging variants in the Lower Midwest was comparable to the number of variants introduced from other regions (Table 1). Variants emerging in this region disseminated only to the Upper Midwest and Northeast, while variants originating from all other regions except the Northeast were introduced into the Lower Midwest (Table 2). A pronounced bidirectional exchange between the Lower and Upper Midwest was observed, indicating strong connectivity between these two interior regions (Table 3; Figures 2D and Figures 3). From a temporal perspective, the Lower Midwest exhibited relatively rapid dispersal dynamics, with locally emerging variants reaching other regions in approximately 1.3 years on average (Table 3; Figures 2D and Figures 3). Dissemination toward the Upper Midwest occurred particularly quickly, with a mean time of approximately 1.3 years (Table 3; Figures 2D and Figures 3).

The Atlantic Seaboard functioned primarily as a source region, exporting variants to most other regions with the exception of the Great Plains, while receiving introductions only from the Northeast and Upper Midwest (Table 3; Figures 2B and Figures 3). This pattern indicates comparatively limited connectivity between the Atlantic Seaboard and the Great Plains. Variants emerging from this region exhibited slower spread dynamics overall, with a mean time to dissemination of approximately 2.6 years (Table 3; Figures 2B and Figures 3).

The Northeast did not function as a major source of PRRSV-2 variants, as most locally emerging variants remained geographically restricted. Export from this region was limited and occurred exclusively toward the Atlantic Seaboard, whereas the Northeast was the only region to receive introductions from all other regions (Table 2). Variants emerging in the Northeast exhibited the shortest average time to dissemination, approximately 1.0 years (Table 3; Figures 2E and Figures 3); however, this estimate is based on only two exportation events and should therefore be interpreted with appropriate caution.

Our findings mirror the previously documented pathways of PRRSV-2 dissemination across the USA, providing further support for the evidence of heterogeneous levels of interconnectivity among major swine-producing regions [56, 57]. Collectively, the time-to-dispersal event analyses (Figure 4) suggest faster dispersal corridors among interior regions (e.g., Great Plains to Upper Midwest and Lower Midwest to Upper Midwest, ∼1.2–2.0 years) and longer for coast to interior pathways (Atlantic Seaboard to interior, ∼2–3 years) (Figure 2 and Figure 3). Dispersal estimates between the Northeast and the Atlantic Seaboard were among the fastest on average, with a mean of approximately one year observed; however, these estimates should be interpreted with appropriate caution due to the low number of observed dispersal events (Figure 2 and Figure 3).

**Figure 4.**
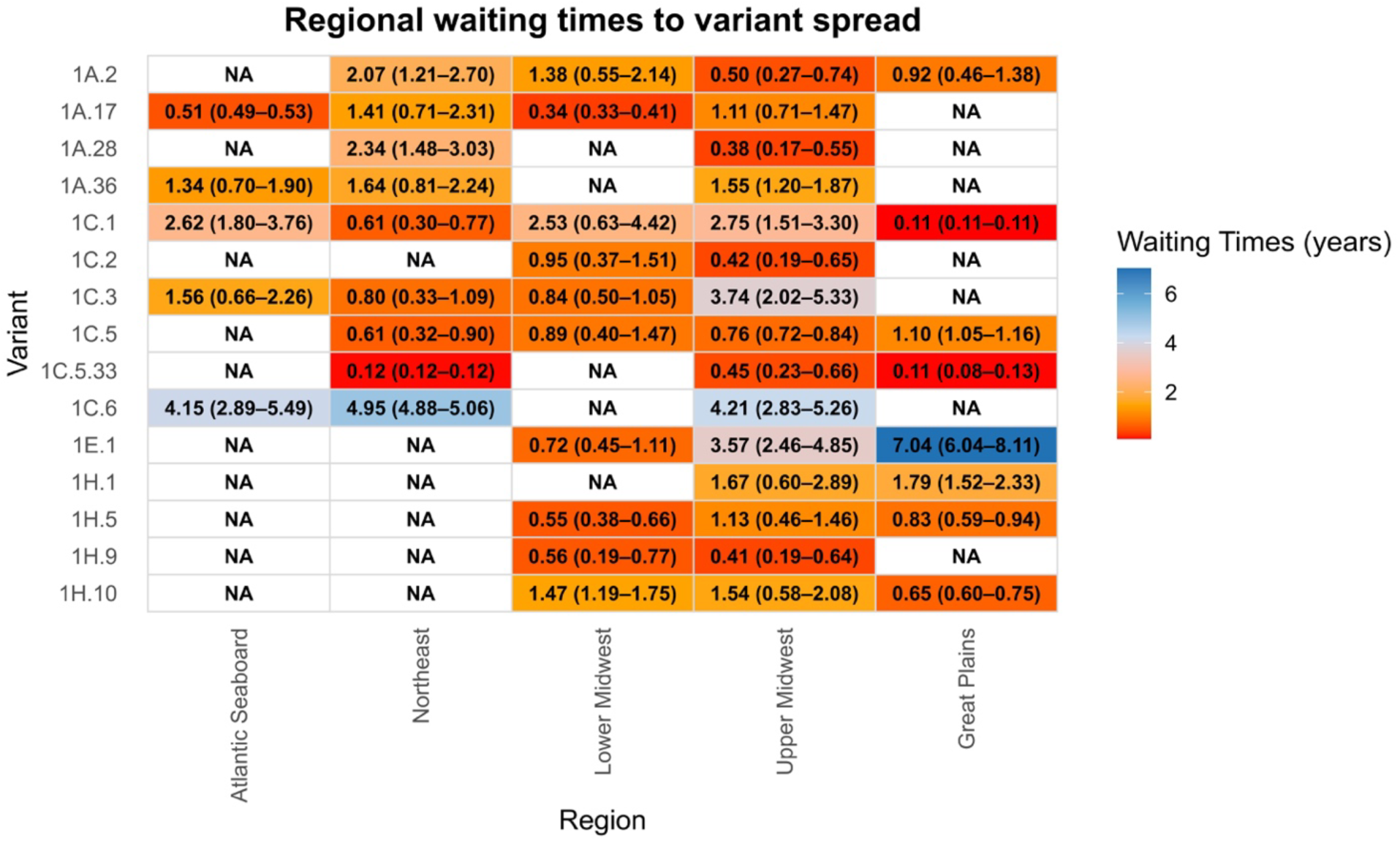
Waiting times of PRRSV-2 variants across USA regions. Persistence was estimated with a structured coalescent approach (PACT). Values represent mean time (in years) with 95% confidence intervals. Warm colors indicate shorter waiting times, meaning variants coalesced more rapidly into distinct regional populations whereas cool colors represent slower dynamics. *NA* indicates regions where the corresponding variant was not detected.

### Structured coalescent inference reveals heterogeneous spread dynamics of PRRSV-2 variants across USA swine-producing regions

Time-to-dispersal event analyses above are independent of genealogical reconstruction and, for this reason, lack the power to capture important nuances of PRRSV-2 dispersal dynamics among distinct regions, such as dispersal events originating from regions other than where a variant first emerged. In addition, dispersal time estimates derived from these approaches depend on the exact dates at which variants sequence was observed, even though variants may have been circulating prior to their detection. To address these limitations, we inferred the time between variant emergence and subsequent dispersal, as well as the spatial trajectories of PRRSV-2 variants, using a phylogeographic framework.

We reconstructed individual time-scaled phylogenies for a total of fifteen variants, each represented by more than 100 sequences and detected in more than one region. This approach allowed us to capture pattern in both the timing and pathways of PRRSV-2 variant dissemination across U.S. regions. The variants analyzed belong to PRRSV-2 sub-lineages 1A (1A.2, 1A.17, 1A.28, 1A.36), 1C (1C.1, 1C.2, 1C.3, 1C.5, 1C.5.33, 1C.6), 1H (1H.1, 1H.5, 1H.9, 1H.10), and 1E (1E.1) (Figure 4 and Supplementary Figure S3-5). The spatial extent of the 15 variants analyzed here varied substantially across variants. Variant 1C.1 was the only variant detected in all regions (Figure 4). Variants 1C.5, 1C.3, 1A.2, and 1A.17 were detected in at least four regions, whereas variants 1A.36, 1C.5.33, 1C.6, 1H.5, 1H.10, and 1E.1 were present in three regions. Finally, variants 1A.28, 1C.2, 1H.1, and 1H.9 were detected in only two regions (Figure 4, Figure 5 and Supplementary Figure S3-5).

**Figure 5.**
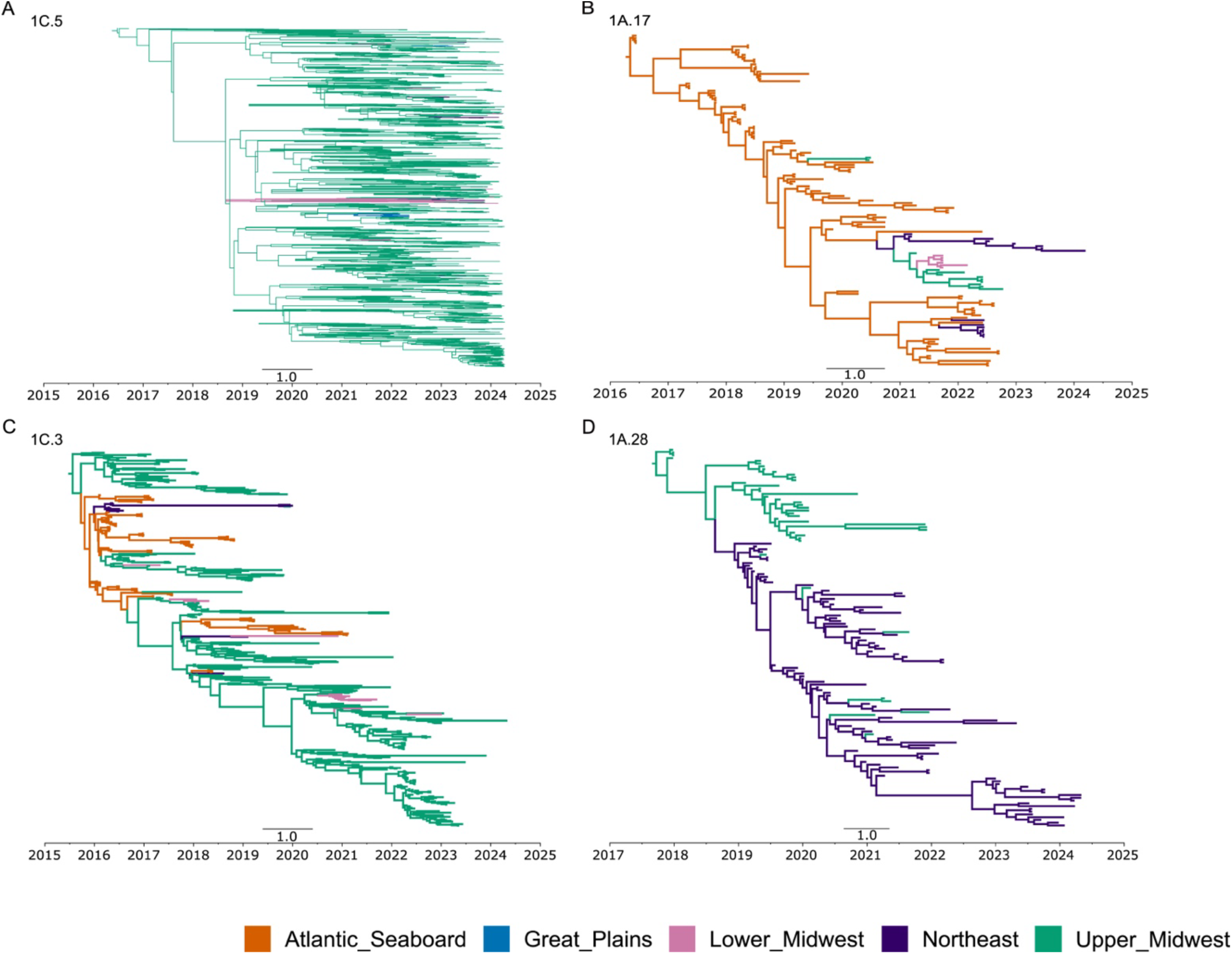
MCMC tree and persistence times of PRRSV-2 variants within sub-lineage 1H. Time-scaled phylogenetic trees (right) and estimated persistence times (left) for four representative variants of sub-lineage 1H across five USA swine-producing regions: Atlantic Seaboard (orange), Great Plains (blue), Lower Midwest (purple), Northeast (pink), and Upper Midwest (green). Phylogenies were reconstructed in BEAST v1.10.4. Panels show: (A) Variant 1C.5, (B) Variant 1A.17, (C) Variant 1C.3, (D) Variant 1A.28.

Analyses of waiting times revealed heterogeneous patterns of inter-regional dispersal across regions. The Great Plains and Lower Midwest exhibited the shortest median estimated waiting times, at approximately 0.87 years (IQR: 0.51–1.28) and 0.86 years (IQR: 0.59–1.27), respectively. The Upper Midwest occupied an intermediate position, with a median waiting time of 1.33 years (IQR: 0.56–2.48), followed by the Northeast, with a median of 1.41 years (IQR: 0.61–2.07) and Atlantic Seaboard showed the longest median waiting time, at 1.45 years (IQR: 0.71–2.35). Overall, these results suggest faster inter-regional spread dynamics in the Great Plains and Lower Midwest, whereas the Upper Midwest, Northeast, and Atlantic Seaboard exhibit comparatively longer waiting times, with substantial overlap in their interquartile ranges.

The waiting times also varied markedly among variants, with substantial heterogeneity both within and across regions. In several cases, dispersal occurred more rapidly in some regions and more slowly in others. Variant 1C.5.33 exhibited the fastest dispersal dynamics overall, with waiting times of approximately 0.11 and 0.12 years in the Northeast and Great Plains, respectively, and 0.45 years in the Upper Midwest. In contrast, variant 1C.6 showed the slowest dispersal dynamics, with waiting times exceeding four years across regions (Figure 4). Some variants displayed highly heterogeneous dispersal patterns depending on the region in which they circulated. For example, variant 1C.3 showed waiting times ranging from approximately 0.77 to 1.56 years in three of the four regions where it circulated but exceeding 3.7 year in the Upper Midwest (Figure 4). An even more pronounced pattern was observed for variant 1E.1, which exhibited the greatest discrepancy in dispersal times across regions, with waiting times of approximately 0.72 years in the Lower Midwest, 3.57 years in the Upper Midwest, and 7.03 years in the Great Plains (Figure 5).

Phylogeographic reconstruction further revealed distinct dispersal routes across variants; four representative phylogeographies displaying various patterns are illustrated in Figure 5. Our results suggest that the great majority of the variants analyzed here emerged in the Upper Midwest, accounting for 8 out of 15 (1A.2, 1A.28, 1C.2, 1C.3, 1C.5, 1C.5.33, 1C.6, and 1H.9) (Figure 5 and Supplementary Figure S3-5). Three variants emerged in the Atlantic Seaboard (1A.17, 1A.36, and 1C.1), three in the Great Plains (1H.1, 1H.5, and 1H.10), and only one in the Lower Midwest (1E.1). Eight variants (1A.2, 1C.1, 1C.2, 1C.5, 1C.5.33, 1H.1, 1H.5, and 1H.9) exhibited a dispersal pattern exclusively from their region of origin to recipient regions, without evidence of onward transmission between recipient regions or reintroduction back to the region of origin (Figure 5 and Supplementary Figure S3-5). In contrast, seven variants (1A.17, 1A.28, 1A.36, 1C.3, 1C.6, 1H.10, and 1E.1) showed dispersal patterns from the region of origin to recipient regions accompanied by subsequent viral exchange among recipient regions, representing alternative dispersal pathways that did not originate exclusively from the original region (Figure 5 and Supplementary Figure S3-5). Reintroduction back to the region of origin was observed for four variants (1A.28, 1C.2, 1C.3, and 1E.1) (Figure 5 and Supplementary Figure S3-5).

## Discussion

Genetic analysis of sequences generated through routine diagnostic testing provides insights of the emergence of novel genetic variants and their evolutionary trajectories and their spatial dynamics over time [18, 78–80]. In this study, we analyzed a comprehensive dataset of PRRSV-2 sequences spanning the five major swine-producing regions of the United States over the last decade (2015-2024). We quantified the genetic diversity of viral populations across regions and estimated the time required for variants to disperse following their emergence, using both a time-to-dispersal event analysis and waiting time analysis.

We observed pronounced heterogeneity in diversity profiles across regions. The Upper Midwest exhibits low diversity despite high richness, where diversity is defined as the probability that two randomly sampled individuals belong do different variants [81]. In contrast, the Lower Midwest is characterized by a more balanced diversity profile, with a more even distribution of variant frequencies.

Indeed, more than 50% of sequences from the Upper Midwest were represented by only four variants (Supplementary Table 2), with variant 1C.5 alone accounting for approximately 30% of all sequences. In contrast, dominant variants in other regions did not reach comparable frequencies. For example, the predominant variant in the Lower Midwest was 1C.2 (11.6% of sequences), in the Northeast 1A.28 (14.3%), in the Atlantic Seaboard 1C.1 (11.6%), and in the Great Plains 1E.1 (21%, Supplementary Table 2). These patterns underscore regional differences in variant frequency distributions across regions and suggest distinct local evolutionary dynamics.

Interestingly, most variants that were prevalent within one region did not appear at similarly high frequencies elsewhere. For instance, variant 1C.5 emerged in the early 2020s and became the predominant variant in the Upper Midwest within approximately two years [58]. Two years after its emergence, 1C.5 was detected in the Lower Midwest, and three years later in both the Great Plains and Northeast; however, it did not establish dominance in any of these regions. Outside the Upper Midwest, 1C.5 reached its highest frequency in the Lower Midwest, where it accounted for approximately 4% of sequences, while in the remaining regions its prevalence did not exceed 1%. Similar patterns were observed for most regionally dominant variants. The exception was variant 1C.2, dominant in the Lower Midwest, which also reached high frequency in the Upper Midwest. Nevertheless, some variants achieved high frequencies in more than one region, including 1C.3 in both the Upper Midwest and Atlantic Seaboard, 1A.2 in the Upper and Lower Midwest, and 1A.36 in the Northeast and Atlantic Seaboard (Supplementary Figure 2).

The underlying factors determining why certain variants become prevalent in specific regions but not others remain poorly understood. Several non-mutually exclusive processes may contribute to these patterns, such as: transmission rates, introduction rates, differential context-dependent fitness, competition with locally circulating variants, and insufficient time for expansion following introduction [82]. In addition, biocontainment efforts after regional introductions may be more aggressive, particularly if a variant has gained notoriety amongst swine health professionals in other regions. The observed heterogeneity in regional diversity profiles further suggests that distinct evolutionary dynamics operate across regions. Importantly, evolutionary mechanisms are acting simultaneously and set of different combinations of their relative contributions may lead to superficially similar diversity patterns [5, 82–84]. Disentangling relative contributions of each mechanisms require additional analyses beyond the scope of this study but represents an important direction for future research.

The Upper Midwest can be considered as a center of diversification for PRRSV-2 variants in the U.S., as it exhibited the greatest variant richness compared with all other regions and accounted for the highest number of locally emerging variants. This region displays unique characteristics that make it an ideal cradle for the emergence of new variants. The Upper Midwest encompasses the U.S. Corn Belt, where a considerable fraction of North American swine production is concentrated, accounting for approximately 48% of USA swine inventory [85]. The high density of animals in this region creates opportunities for more intense viral circulation amongst herds. In addition, the frequent movement of growing pigs into the Upper Midwest facilitates the introduction of viruses from other regions [34, 50–52, 55]. Together, these factors increase the opportunity for co-infections with distinct variants, which is a prerequisite for recombination events that sometimes result in novel viral variants [5, 27, 86–91].

We also observed asymmetry in the dissemination of PRRSV-2 variants among the five regions: the Upper Midwest serves as a central node in the transmission network, serving simultaneously as both a source and a recipient region, while the Atlantic Seaboard exhibited asymmetric connectivity, appearing more isolated from a recipient standpoint, receiving fewer introductions from other regions, while simultaneously serving as a source of introductions into other regions, corroborating previously demonstrated patterns. [51, 56]. Furthermore, dispersal times varied considerably across region pairs, with variants from other regions tending to reach the Lower Midwest more rapidly than other destinations, while introductions from the Atlantic Seaboard into the other regions occurred more slowly. Together, these findings expand our understanding of the spatiotemporal dynamics of PRRSV-2 in the United States, offering a broader view of how interregional connectivity influences the timing and patterns of viral dissemination. However, dispersal times are unlikely to be uniform across variants and are shaped by a combination of epidemiological, demographic, evolutionary, and ecological dynamics.

Both methodological approaches utilized in this paper (the time-to-dispersal event analysis and waiting time analysis) presented complementary strengths and limitations. The time-to-dispersal event analysis allowed us to estimate dispersal intervals for a large number of variants, including those with limited sequence representation, which would be more difficult to address using ancestry-based reconstruction models. However, this approach inevitably underestimates introduction times, since variants may circulate undetected for some time before reaching detectable levels; our definition of emergence as the fifth report, a decision made to reduce noise, likely further contributes to this bias. In addition, the time-to-dispersal event analysis does not provide a broader view of alternative routes by which variants are introduced into different locations. For example, exchanges of variants from the Northeast to the Upper Midwest were not detected in the time to dispersal event-based estimates, since this approach focuses exclusively on locally emerging variants and their subsequent dispersal to new regions. In contrast, coalescent-based phylogenies suggest that migration events have occurred from the Northeast to the Upper Midwest, serving as an alternative pathway for certain variants. However, this was not detected very frequently; the only one such case was variant 1A.17 (Figure 5B), which appears to have initially emerged in the Atlantic Seaboard, was subsequently introduced into the Northeast, and ultimately reached the Upper Midwest. The structured coalescent framework provided more detailed insights into persistence within regions, potential routes of spread, and independent introduction events. Yet, its application was constrained to variants with sufficient sequence data, excluding variants with smaller sample sizes.

Despite some variation between the two analyses, both approaches consistently identified the Lower Midwest as having quick inward and outward dispersal times, the Upper Midwest as occupying an intermediate position, and the Atlantic Seaboard as exhibiting comparatively slower inter-regional dynamics. The main discrepancies were observed for the Great Plains and the Northeast, likely reflecting differences in how each analysis captures dispersal process, or alternatively resulting from differences in the underlying datasets, as the waiting-time analysis relies on a subset of variants with sufficient number of sequences, whereas the time-to-spread framework incorporates a more comprehensive dataset that includes lower-frequency variants. Together, these results underscore the complementary nature of the two approaches and highlight the importance of integrating multiple analytical frameworks to capture the full complexity of PRRSV-2 spread across production regions.

Although the number of sequences analyzed varied among regions, rarefaction analyses suggest that our sampling effort was may have been sufficient to capture most of the regional diversity of variants, and that the number of sequences is broadly proportional to swine herd density across the United States [92]. Sequences classified as “unclassified variants” were not included in the diversity analyses, as they represent poorly resolved phylogenetic groups. Consequently, the true diversity in each region may be higher than reported here, and this limits our overall understanding of spatial and temporal dynamics.

In summary, our findings are valuable for surveillance of PRRSV-2 dispersal across U.S. swine-producing regions, as they highlight faster transmission corridors across which viral variants spread more rapidly. These insights can support the implementation of early warning systems, improving communication among swine production systems and enabling more timely and targeted responses to emerging variants. Such information may also inform management strategies by introducing a temporal focus to intervention efforts and facilitating the deployment of stricter biosecurity and biocontainment measures in regions that are highly connected and tend to disseminate variants more rapidly.

These results can further inform timely containment measures, particularly in highly connected regions where the risk of rapid dissemination is greatest. In addition, the estimated dispersal times provide useful reference points for calibrating epidemiological models, offering more accurate parameters on the pace of variant spread and supporting the development of robust predictive frameworks to assess the risk of new introductions. However, these estimates should not be regarded as absolute values, but rather as general indicators of the temporal lag between emergence and dispersal. The dispersal time of individual variants may be shaped by intrinsic viral biological characteristics, host population demographic dynamics, and varying levels of connectivity among production regions, among other factors.

## Author Contributions

Conceptualization: J.P.H.S., and K.V.; methodology: J.P.H.S, and K.V.; Investigation: J.P.H.S., and K.V.; database curation: J.P.H.S, M.K. and C.A.C.; formal analysis: J.P.H.S., and K.V.; original draft writing: J.P.H.S.; review and editing: J.P.H.S., K.V., I.A.D.P., M.K., and C.A.C.; Supervision: K.V.; Funding acquisition: K.V. All authors have read and agreed to the published version of the manuscript.

## Acknowledgments

The authors would like to thank the Morrison Swine Health Monitoring Project (MSHMP) participants and the University of Minnesota Veterinary Diagnostic Laboratory for sharing its PRRSV-2 genetic sequences.

